## Supplemental Figures and Tables for "Long-distance dispersal of pigeons and doves generated new ecological opportunities for host-switching and adaptive radiation by their parasites"

### Supplementary figures and tables

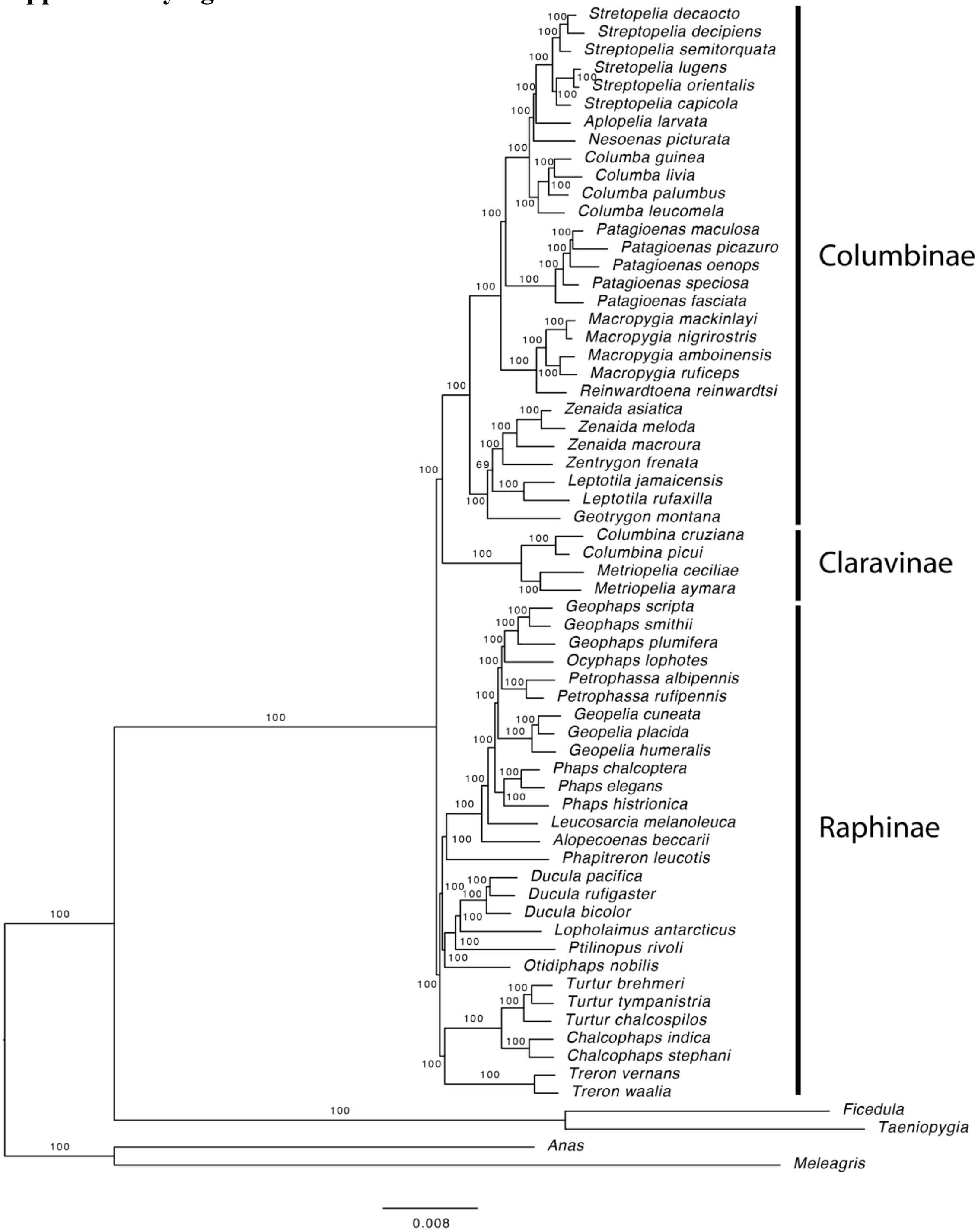


**Figure S1**. Maximum-likelihood tree showing the relationships of dove species based on the concatenation of 6,363 single-copy orthologous gene sequence alignments. The tree was calculated using RAxML and support was calculated from bootstrap replicates. Numbers at nodes indicate support as percent of 100 bootstrap replicates that also recovered the same node. Vertical bars and names indicate subfamilies.

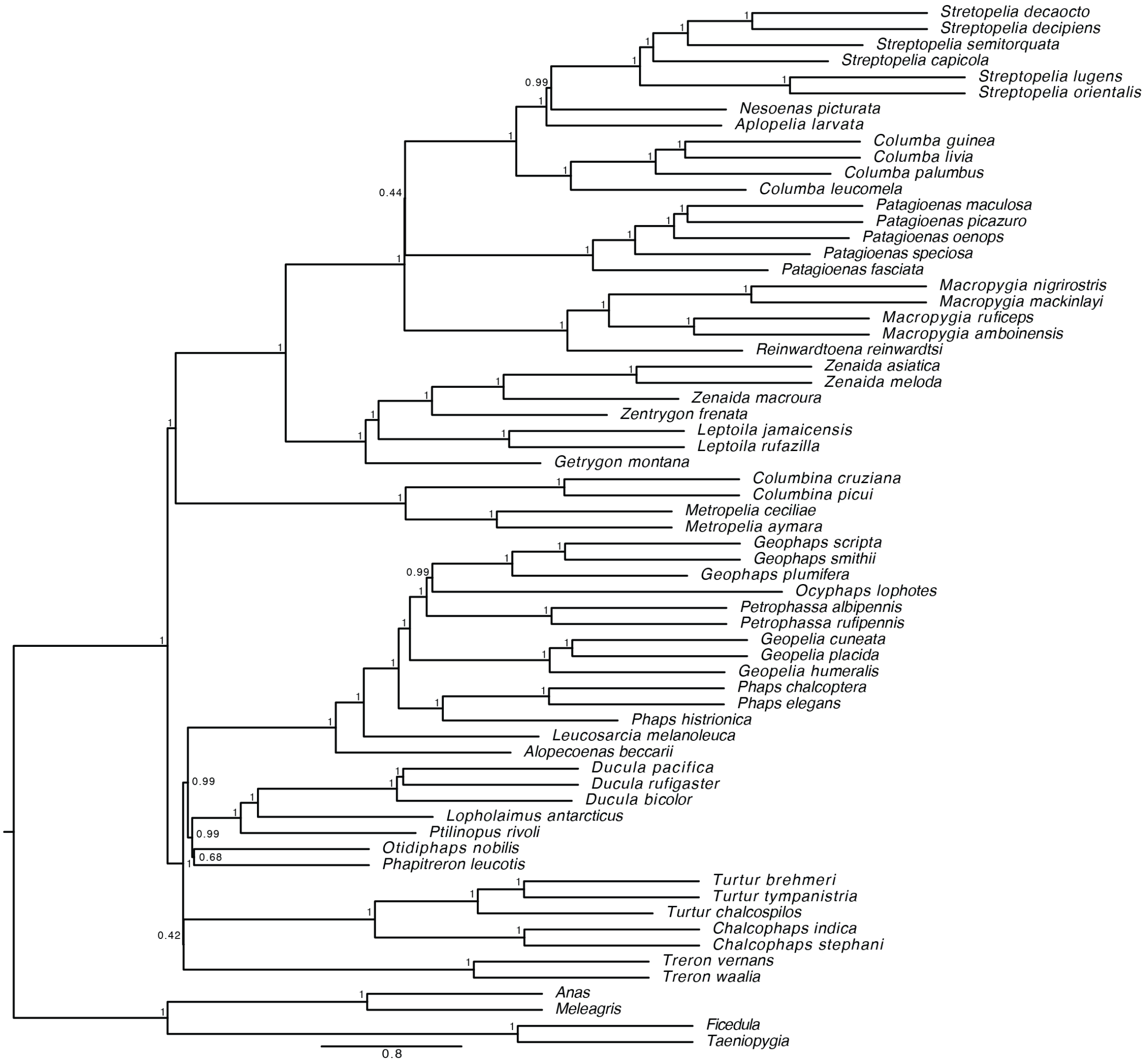


**Figure S2**. Coalescent tree showing relationships of dove species based on 6,363 single-copy ortholog gene trees. Tree was calculated using ASTRAL from gene trees calculated using RAxML and numbers at nodes indicate support based on local probabilities.

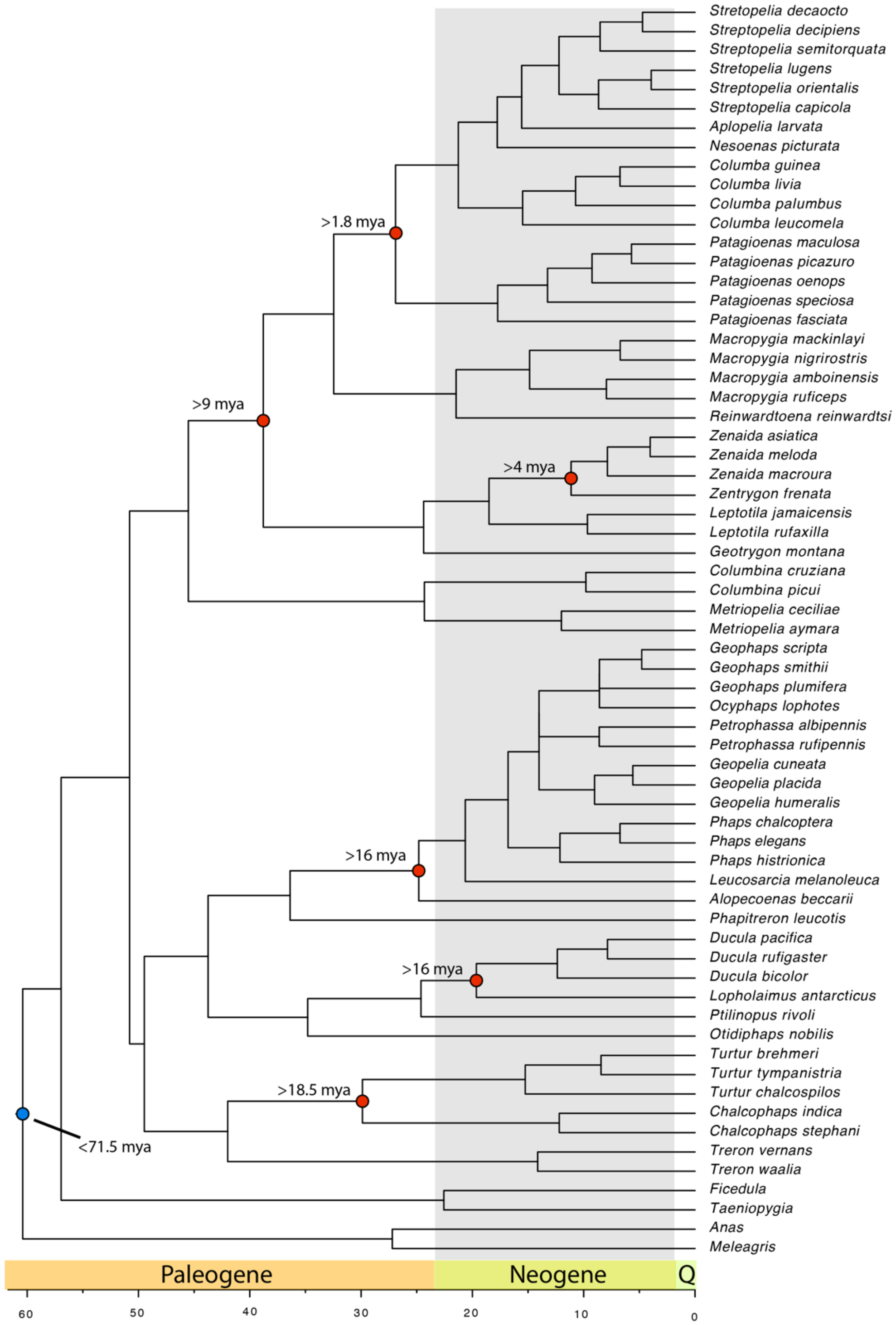


**Figure S3**. Time calibrated maximum-likelihood tree showing the relationships of dove species based on the concatenation of 6,363 single-copy orthologous gene sequence alignments. Tree calibrated with a root of 74.5 mya.

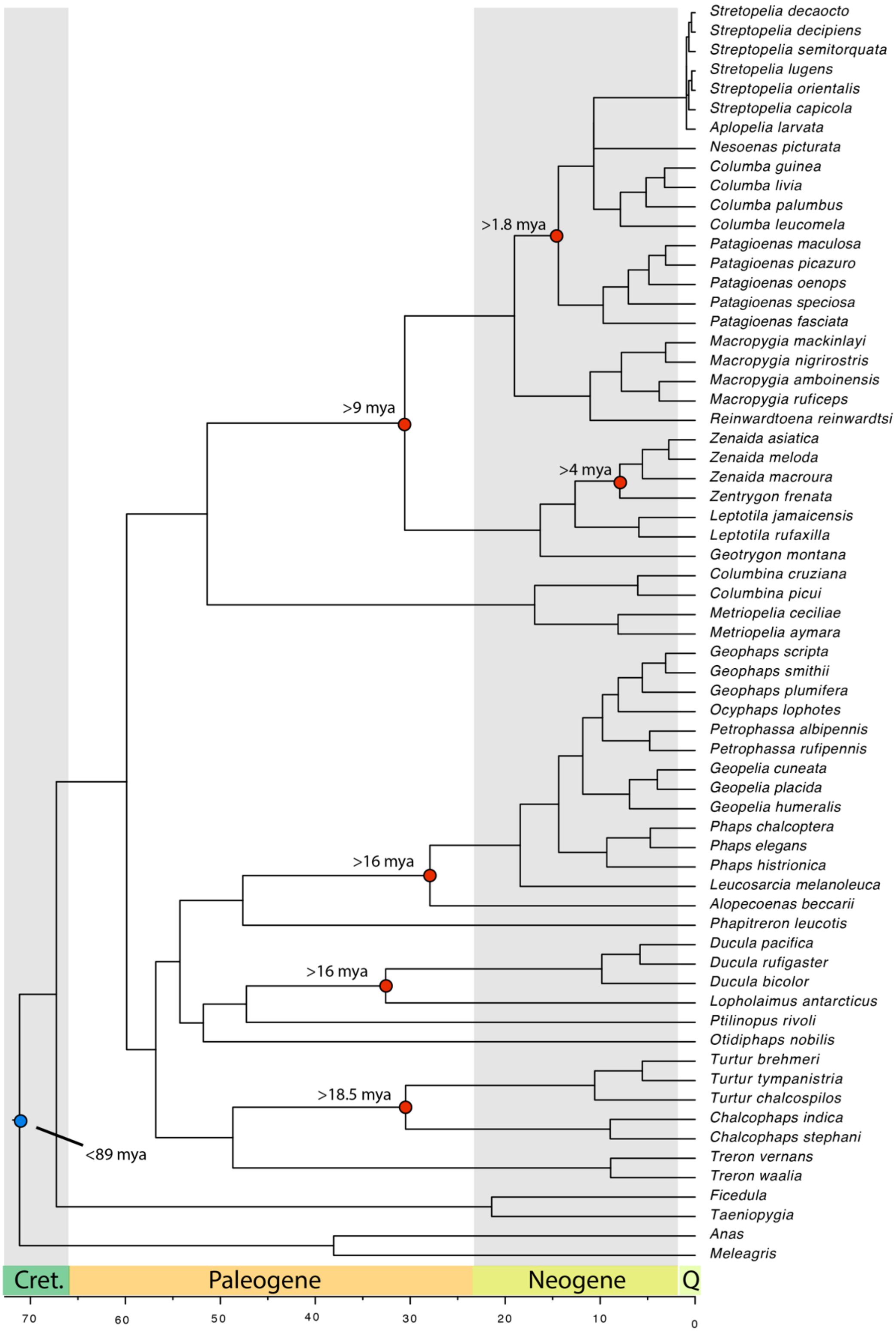


**Figure S4**. Time calibrated maximum-likelihood tree showing the relationships of dove species based on the concatenation of 6,363 single-copy orthologous gene sequence alignments. Tree calibrated with a root of 89 mya.

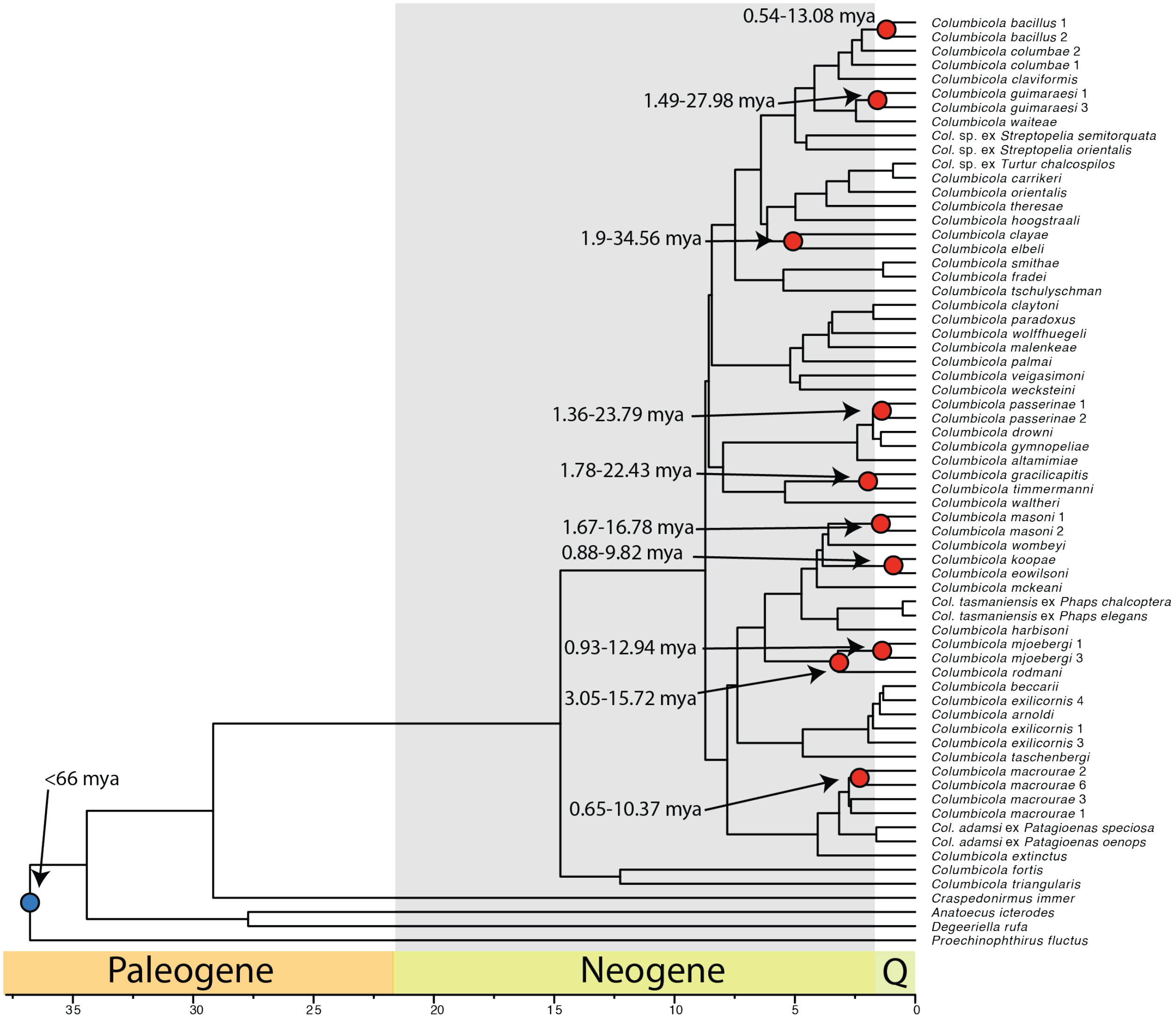


**Figure S5.** Time calibrated maximum-likelihood tree showing the relationships of wing louse species based on the concatenation of 977 single-copy orthologous gene sequence alignments. Tree calibrated with a root of 66 mya and candidate cospeciation events identified from divergence estimates in the dove tree in figure S3.

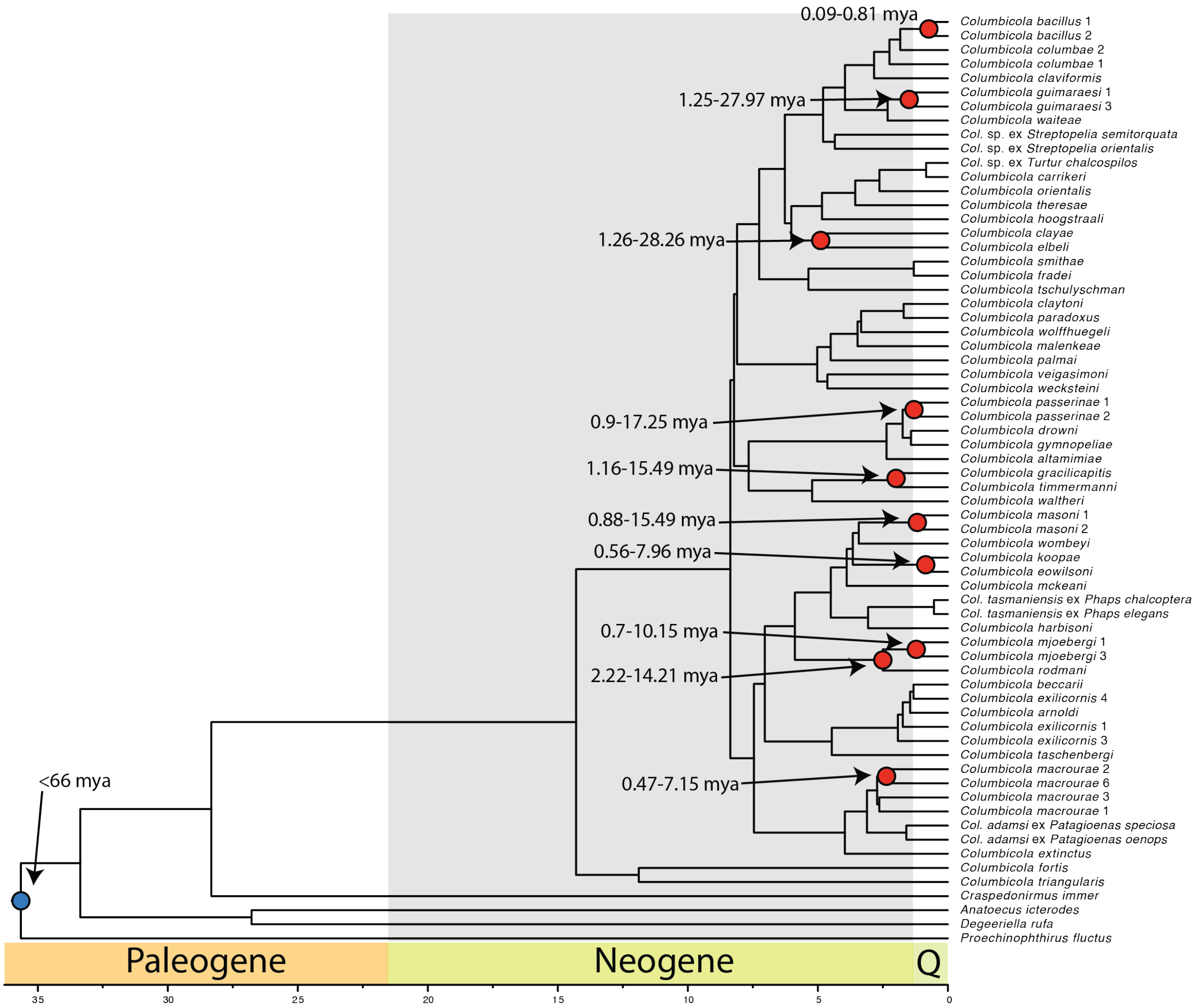


**Figure S6.** Time calibrated maximum-likelihood tree showing the relationships of wing louse species based on the concatenation of 977 single-copy orthologous gene sequence alignments. Tree calibrated with a root of 66 mya and candidate cospeciation events identified from divergence estimates in the dove tree in figure S4.

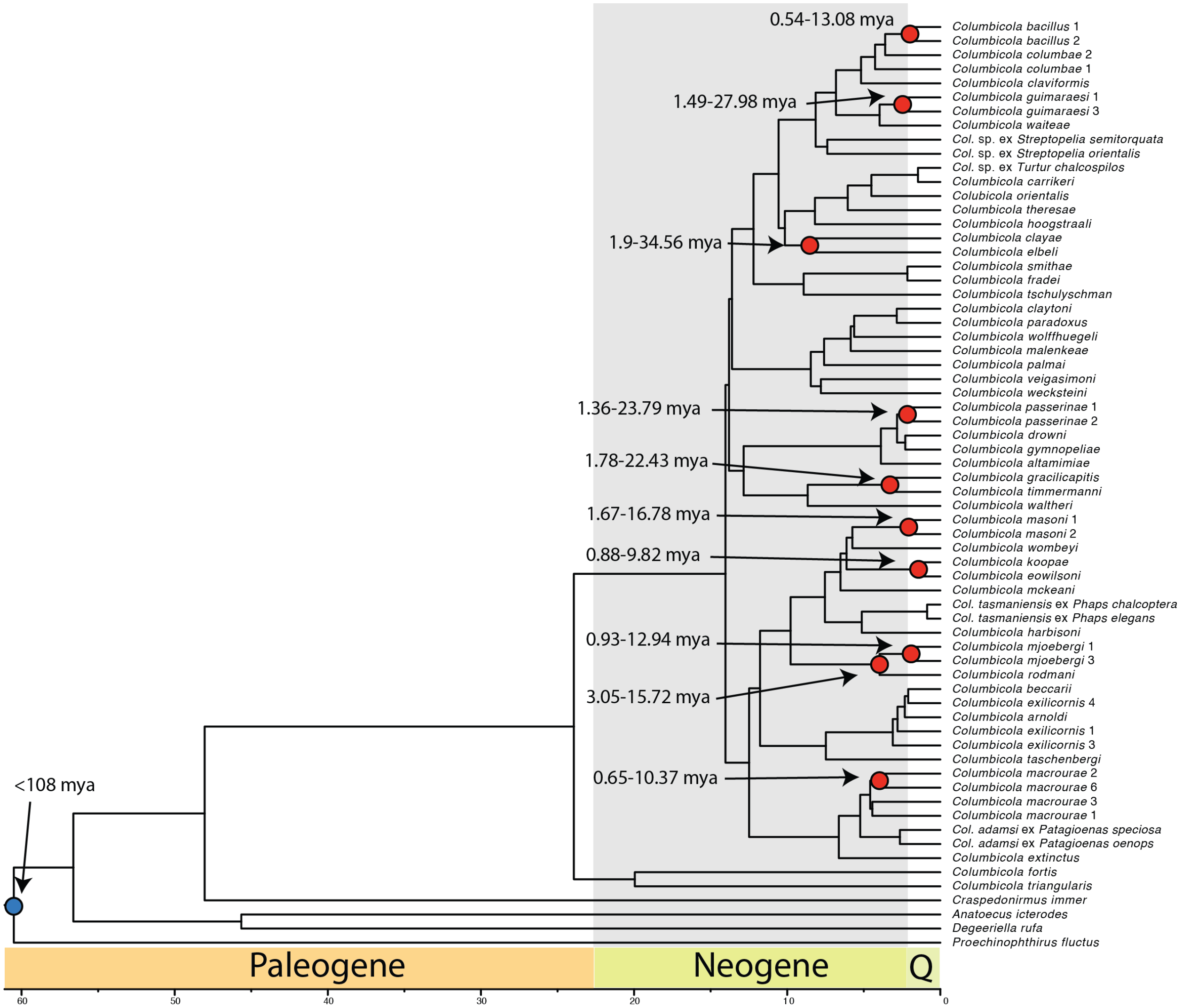


**Figure S7.** Time calibrated maximum-likelihood tree showing the relationships of wing louse species based on the concatenation of 977 single-copy orthologous gene sequence alignments. Tree calibrated with a root of 108 mya and candidate cospeciation events identified from divergence estimates in the dove tree in figure S3.

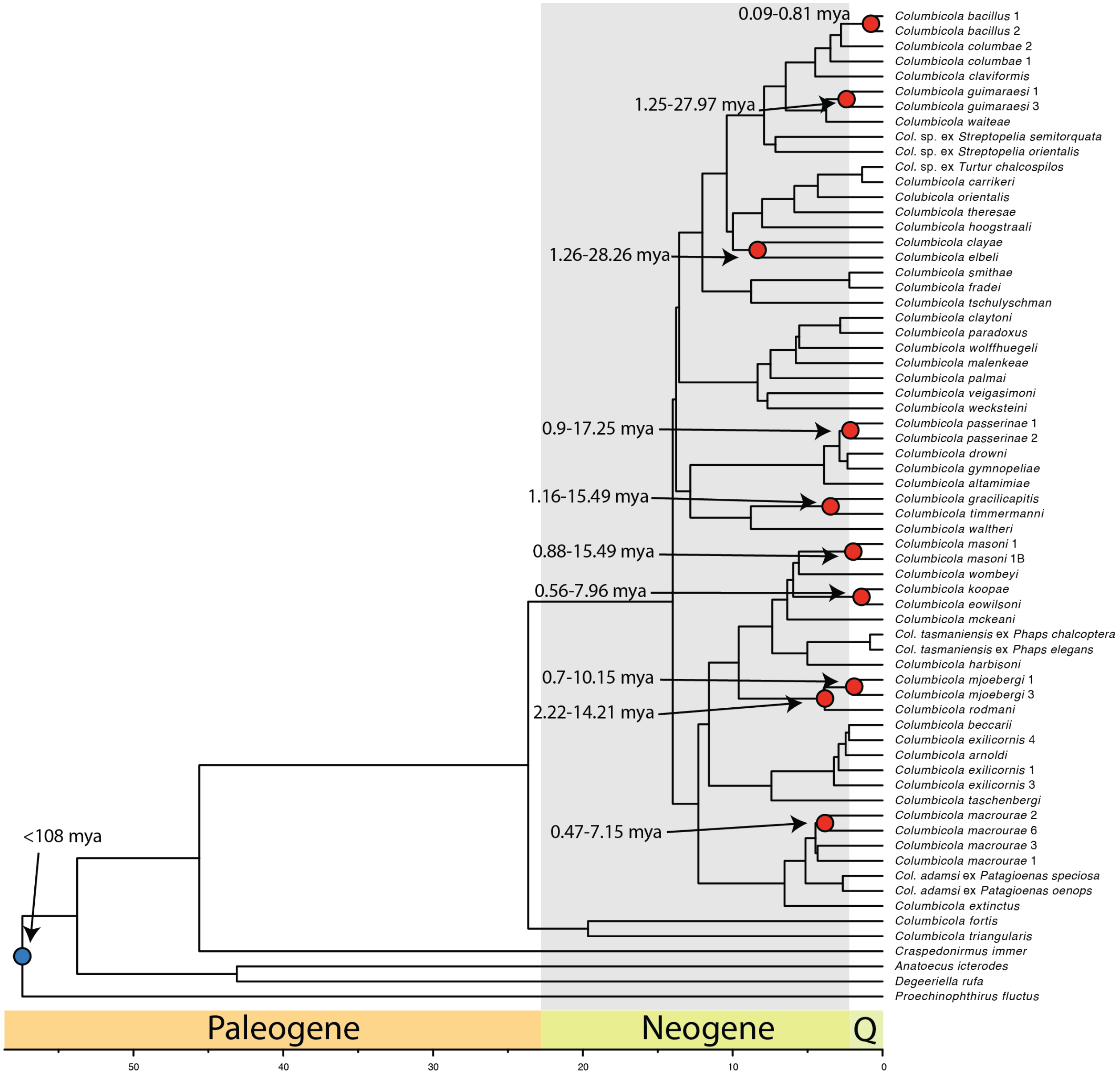


**Figure S8.** Time calibrated maximum-likelihood tree showing the relationships of wing louse species based on the concatenation of 977 single-copy orthologous gene sequence alignments. Tree calibrated with a root of 108 mya and candidate cospeciation events identified from divergence estimates in the dove tree in figure S4.

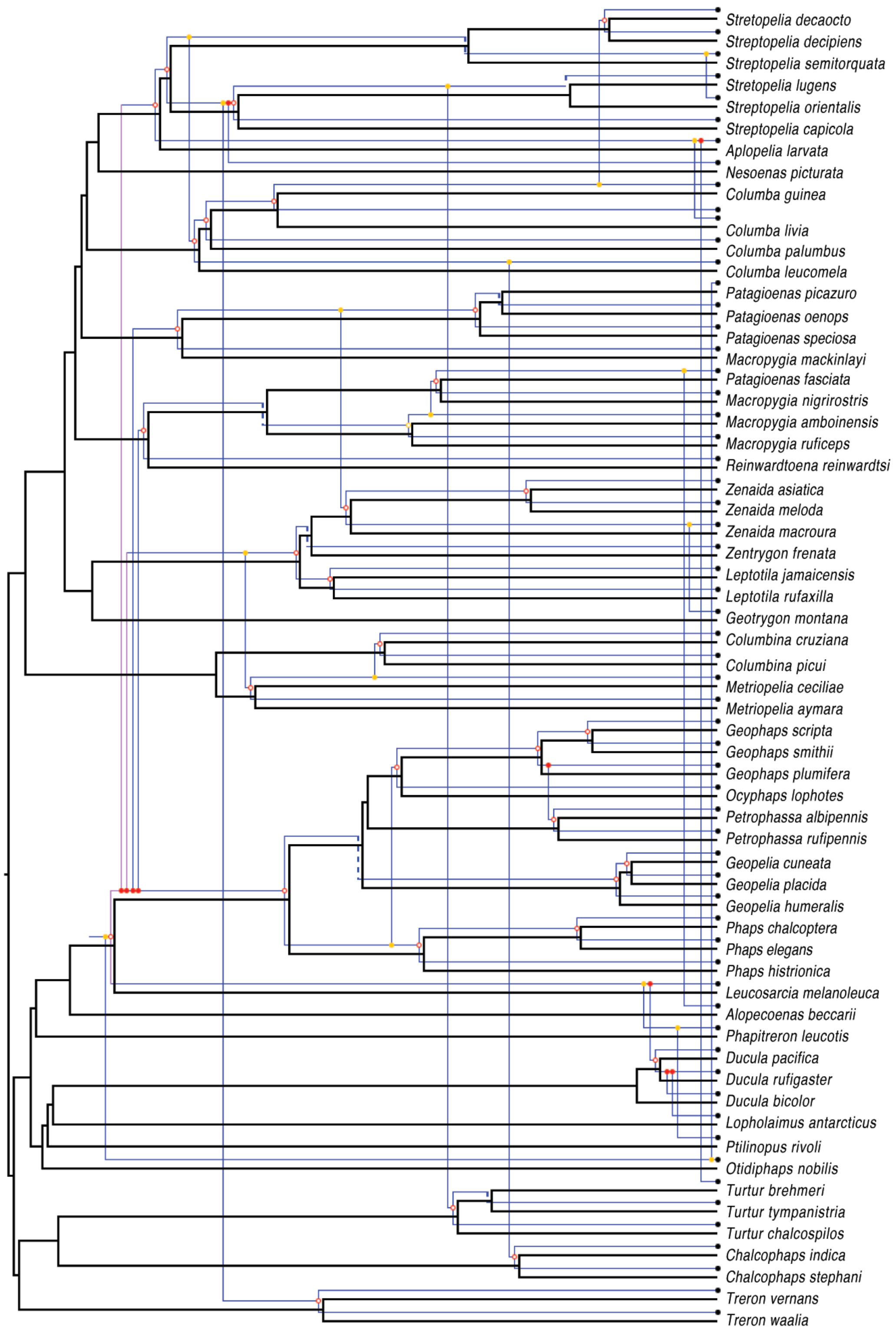


**Figure S9.** JANE cophylogeny reconstruction of doves and wing lice with an Australasian origin of wing lice. Host phylogeny is black and parasite phylogeny is blue.

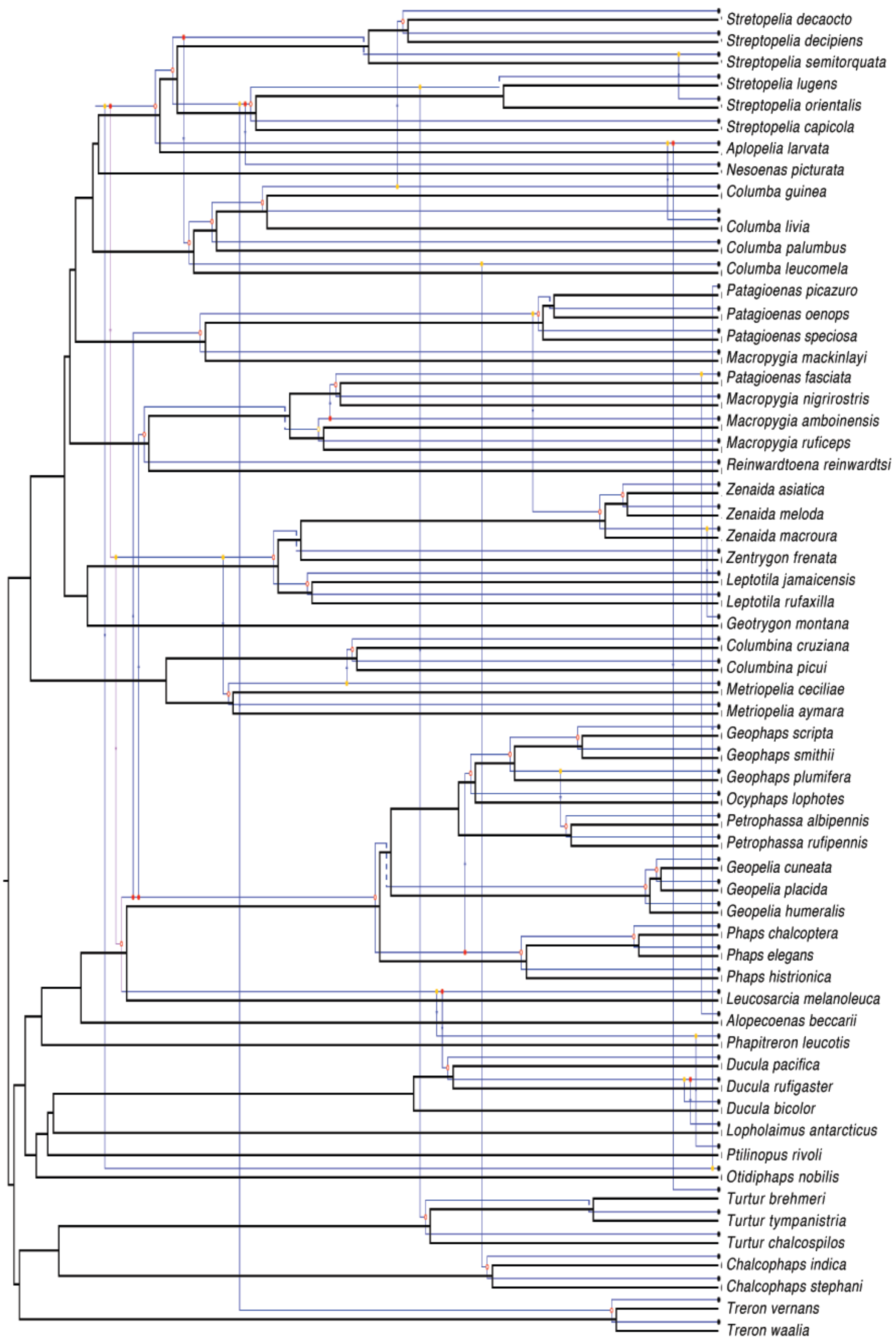


**Figure S10.** JANE cophylogeny reconstruction of doves and wing lice with an African origin of wing lice. Host phylogeny is black and parasite phylogeny is blue.

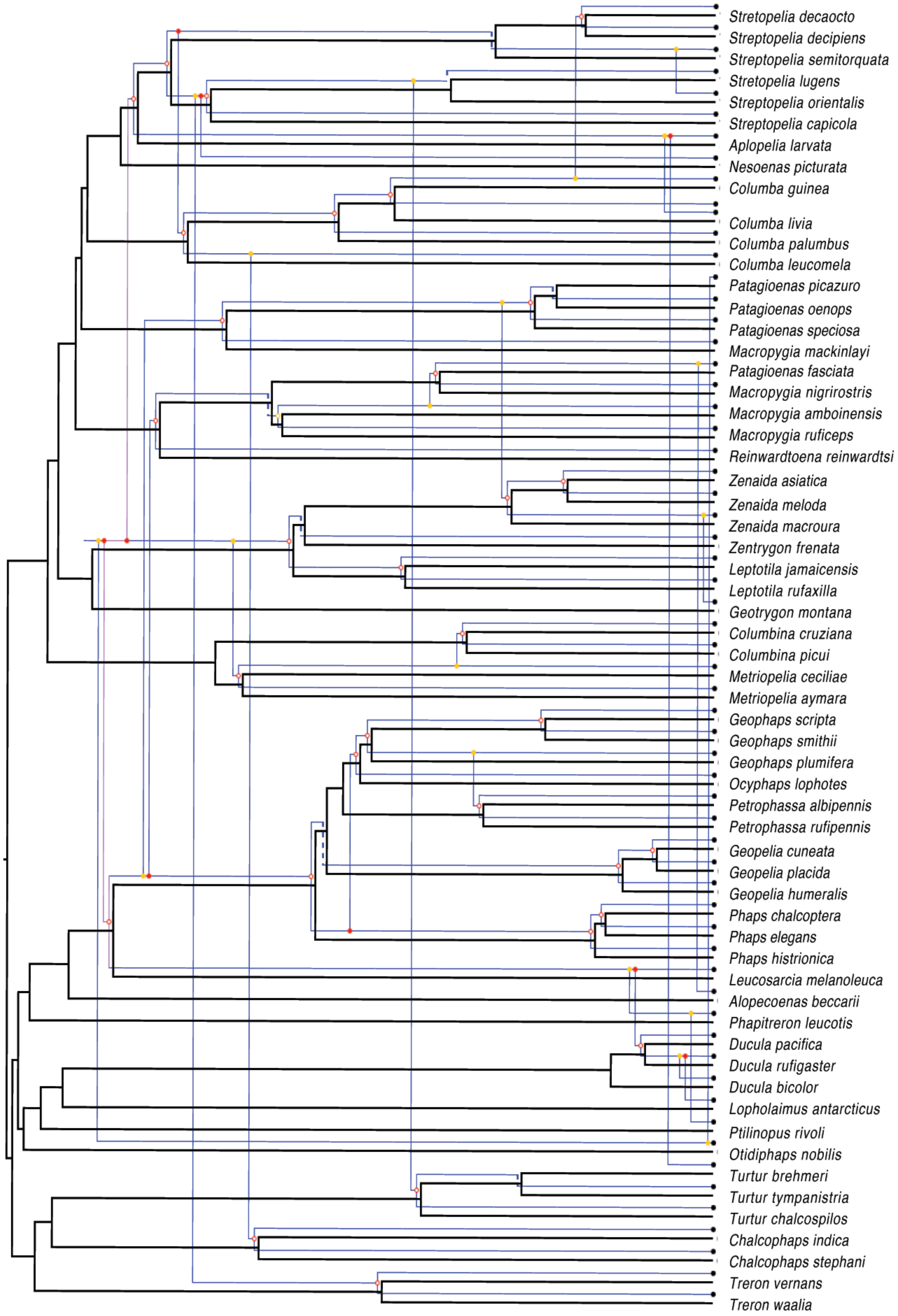


**Figure S11.** JANE cophylogeny reconstruction of doves and wing lice with a New World origin of wing lice. Host phylogeny is black and parasite phylogeny is blue.

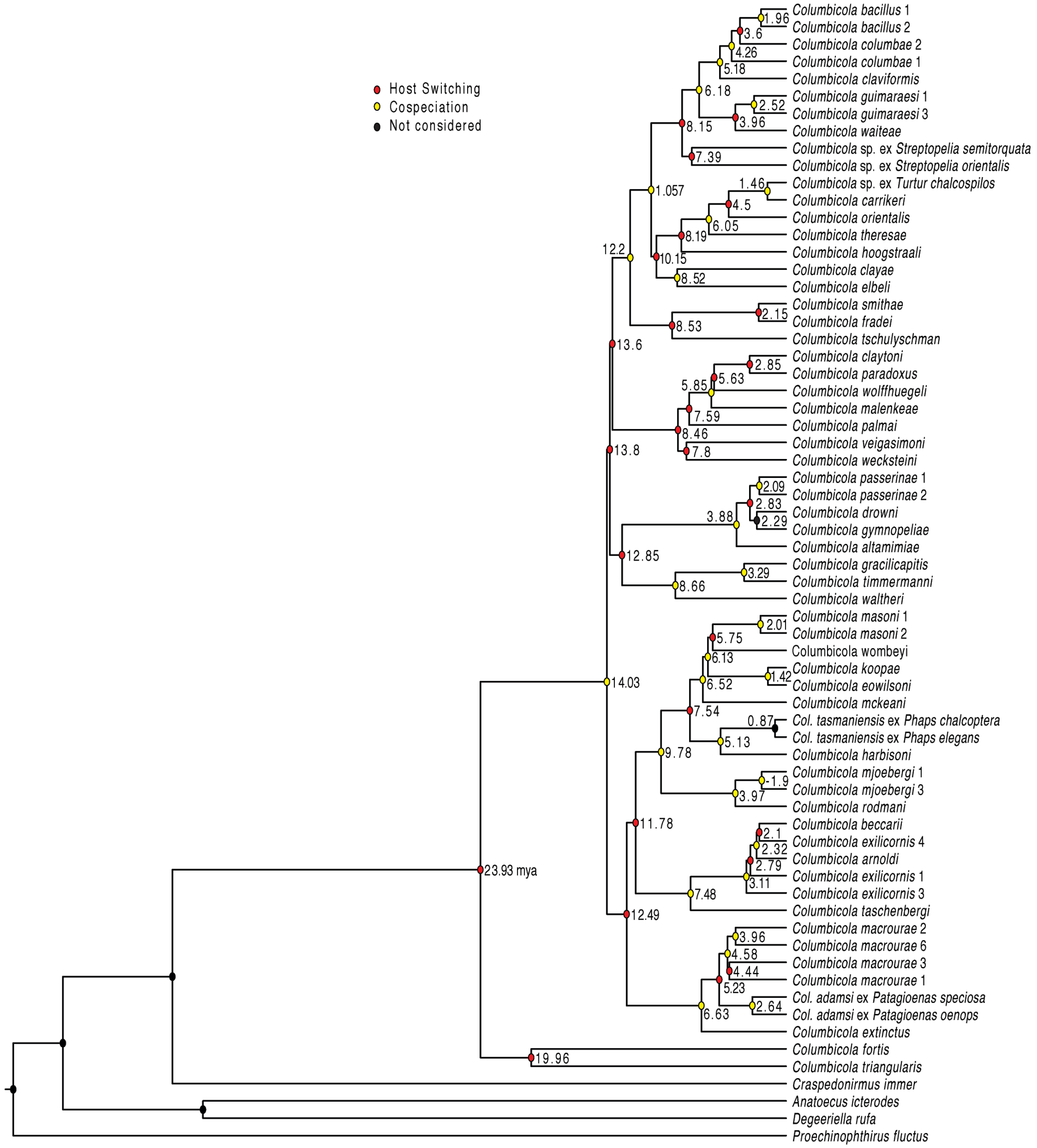


**Figure S12.** Time calibrated maximum-likelihood tree showing the relationships of wing louse species based on the concatenation of 977 single-copy orthologous gene sequence alignments. Tree calibrated with a root of 108 mya and candidate cospeciation events identified from divergence estimates in the dove tree in figure S3. Node labeles indicate node age in millions of years. Node color indicates cospeciation or host-switching events, based on an Australasian origin of wing lice described in figure S9.

**Table S1.** Calibration points used to determine species divergence times for dove and wing louse species.

| Tree | Constraint | Age in mya | Data type | Source |
|---|---|---|---|---|
| host | *Turtur* and *Chalcophaps* | 18.5 | fossil | [6] |
| host | *Lopholaimus* and *Ducula* | 16 | fossil | [7] |
| host | *Alopecoenas* and phabines | 20 | fossil | [8] |
| host | *Zenaida* and *Zentrygon* | 4 | fossil | [9] |
| host | *Geotrygon* and NWGD | 9 | fossil | DWS per. comm. |
| host | *Patagioenas* and *Columba* | 1.8 | fossil | [10] |
| host | root | 71.5 | inference | [11] |
| host | root | 89 | inference | [12] |
| parasite | *C. macrourae* 2 and *C. macrourae* 6* | variable^ | inference | this study |
| parasite | *C. rodmani* and *C. mjoebergi* | variable^ | inference | this study |
| parasite | *C. koopae* and *C. eowilsoni* | variable^ | inference | this study |
| parasite | *C. masoni* 1 and *C. masoni* 2* | variable^ | inference | this study |
| parasite | *C. guimaraesi* 1 and *C. guimaraesi* 3* | variable^ | inference | this study |
| parasite | *C. bacillus* 1 and *C. bacillus* 2* | variable^ | inference | this study |
| parasite | *C. clayae* and *C. elbeli* | variable^ | inference | this study |
| parasite | *C. gracilicapitis* and *C. timmermanni* | variable^ | inference | this study |
| parasite | *C. passerinae* 1 and *C. passerinae* 2* | variable^ | inference | this study |
| parasite | root | 108 | inference | [13] |
| parasite | root | 66 | inference | [14] |

NWGD = New World Ground-doves, DWS = David Steadman, *C.* = *Columbicola*. * designates divergent lineages of lice that were recognized and designated by numbers following Boyd et al. [16], ^ designates that the age of calibration changes based on alternative host tree calibrations (figures S5-S8).

**Table S2.** Comparison of host and parasite trees under different coevolutionary scenarios

| Host diversification | Time zones | Possible origins | Expected cost M | Expected cost SD | Observed cost | Cospecation events | H-S triggered event |
|---|---|---|---|---|---|---|---|
| not used | not used | 1 | 107 | 2 | 61 | 35 | 25 |
| 51 mya | 3 | 3 | 107 | 2 | 63 | 32 | 28 |
| 60 mya | 3 | 4 | 107 | 3 | 63 | 32 | 28 |

M = mean, SD = standard deviation. H-S triggered events = parasite speciation triggered by a host switch. The "possible origins" column denotes the sum of mutually exclusive events.

### Supplementary methods

**Sequencing and quality control**: Kapa Library Preparation kit was used to prepare gDNA libraries. Four libraries were sequenced on one lane on the Illumina HiSeq 2500 using the HiSeq SBS Sequence Kit v2 for 2x160 cycles. The remaining libraries received 6-base barcodes and were pooled into groups of two for sequencing on the same instrument for 2x251 cycles. Resulting sequence reads were trimmed to remove bases with a phred score <30 from the 5' end using Trimmomatic PE (v0.22) [1].

**Identification of within clade paralogs**: A total of 6,923 genes were identified as single-copy orthologs using OrthoDB v8 [2]. The *Columba livia* ortholog set was indexed using Bowtie2 (v2.1.0) [3] and genome sequence reads from *Columba rupestris* were mapped onto the orthologs. Average sequencing depth of each ortholog was calculated using SAMtools and BEDtools (v0.1.19) [4]. Summary statistics were calculated for sequencing depth across orthologs and a z-score of 2.5 standard deviations was established to identify outliers. Those genes with a mean sequencing depth of $-2.5<Z>2.5$ were removed from the ortholog set.

**Ortholog assembly**: Sequence reads from each library were assembled using the *C. livia* orthologs as a reference sequence using Bowtie2 (v2.3.4.1) [3]. Results of read to gene alignments were converted to a binary alignment map and sorted using SAMtools (v1.7). Finally, consensus sequences were created using SAMtools and vcfutils.pl (vcf2fq) by taking the consensus base at each position, yielding ortholog sequences that could be used for phylogenetics. We conducted a final check of overall sequencing coverage for all genes in all taxa to ensure there were no high-coverage sequences, which might signify repeat elements.

**Model fitting and quality control in supermatrix generation**: To control for substantial variation in base composition in the concatenated ML tree, we estimated individual GTR+Γ rate parameters for all codon positions across all genes and removed any position that had a rate parameter that deviated more than 10 standard deviations from the mean. We then clustered the codon positions based upon GTR+Γ parameters using k-means clustering to generate partitions for the ML estimates that would group codon partitions with similar rate matrices into the same partition. After varying the number of partitions to be between 2 and 30, we found that seven partitions explained more than 75% of the total variance of the clustering.

**Dove MCMCtree parameters**: Bayesian MCMC searches were conducted for 10 million generations, sampling every 100 generations with the following parameters: 71.5 calibration (alpha = 0.19563, and two parameters for rgene_gamma (1 12.01) and sigma2_gamma (1 10)); 89 calibration (alpha = 0.19, rgene_gamma (1 15.49) sigma_2gamma (1 10)) as estimated from baseml. The first 50,000 trees were excluded as burn-in for each run. Implemented in PAML (v4.9) [5].

**Louse MCMCtree parameters**: Bayesian MCMC searches were conducted for 5 million generations, sampling every 50 generations with the following parameters: 108 calibration (alpha = 0.275, and two parameters for rgene_gamma (680 1000 1.0 0) and sigma2_gamma (0.77 100 1.0)); 66 calibration (alpha = 0.275, rgene_gamma (1115.7 1000 1.0) sigma_2gamma (0.01 100 1.0)) as estimated from baseml. The first 50,000 trees were excluded as burn-in for each run.

### Supplementary data

The following is included in the figshare database 10.6084/m9.figshare.12730337: raw coding sequences, gene trees, ASTRAL tree, concatenation alignment, concatenation partition, concatenation RAxML-bipartition tree, base frequency by codon image, MCMC trees, Jane input files, sample collection data, NCBI-SRA identifiers, and list of sample collections identifiers.
